# Ferritin Nanoparticle-Based Indirect ELISA for Immunodominant Region Screening and B-cell Epitope Validation of African Swine Fever Virus

**DOI:** 10.64898/2025.12.28.696791

**Authors:** Han Ding, Liubao Du, Suqiu Wang, Junyao Zhong, Jingru Zhang, Jie Li, Lifeng Zhang, Xingqi Zou, Hanchun Yang, Nianzhi Zhang

## Abstract

African swine fever virus (ASFV) poses a severe threat to the global pig industry, creating an urgent need for effective vaccines and reliable diagnostic tools. While numerous ASFV B-cell epitopes have been reported based on murine-derived monoclonal antibodies (mAbs), such epitopes may not fully represent the antibody targets recognized in naturally infected pigs. Therefore, we need an approach to screen immunodominant regions by evaluating the reactivity of pre-defined epitopes with ASFV-infected pigs’ sera. In this study, we developed an indirect enzyme-linked immunosorbent assay (iELISA) based on recombinant epitope-ferritin nanoparticles (EFNs), as a platform for immunodominant region screening and epitope validation. This assay, hereafter referred to as EFNs-iELISA, shows an approximately twofold improvement in sensitivity compared with synthetic peptide-based ELISA for linear B-cell epitopes, and improved discrimination over GST-His₆-peptide and BSA-conjugated peptide formats. Using EFNs-iELISA, we evaluated 22 known murine-derived B-cell epitopes and 17 predicted epitopes from ASFV proteins CD2v, P30, P54, and P72 with ASFV-positive and negative porcine sera. According to the calculated cut-off threshold, tested epitopes were classified into different groups with positive or negative reactivity. Our analysis confirmed that among 22 known murine-derived ASFV B-cell epitopes, 16 (72%) also showed positive reactions with porcine-derived sera. What’s more, we also screened 7 new porcine-specific immunoreactive regions on these proteins. The EFNs-iELISA method provides a practical tool for screening immunodominant regions of ASFV, with potential applications in vaccine and diagnostic research.

## 1. Introduction

African swine fever (ASF) is a highly contagious and deadly disease caused by the African swine fever virus (ASFV)[1]. It is characterised by high fever and subcutaneous hemorrhaging. ASF was first reported in Kenya in 1921. Over the following century, it spread to Cuba, Brazil, Spain, France, Italy, the United States, and Poland, causing significant damage to the global pig farming industry[2]. The initial ASF outbreak in China occurred in August 2018, resulting in economic losses exceeding 10 billion US dollars[3].

ASFV genome is approximately 170 kb-190 kb in length, and encodes more than 160 proteins[4]. The ASFV EP402R gene is a critical protein found on the outer membrane of the virus[5]. It’s a protein homolog of the T-lymphocyte surface adhesion factor CD2, which allows it to mediate the hemadsorption phenomenon[6]. In addition, CD2v can induce a cross-protective immune response against ASFV infection, and the antibodies it elicits can neutralize the virus[7]. Because of these qualities, CD2v is widely used in ASFV vaccine development[8]. P72 is a major structural component of the viral capsid encoded by the ASFV B646L gene[9]. It is not only essential for viral adsorption and cell invasion but also induces high levels of specific antibodies in hosts[10]. As a highly immunogenic target, P72 has become a key biomarker for ASFV diagnosis and serological identification[11]. P30 is a structural protein of the viral inner membrane, encoded by the ASFV CP204L gene[12]. It induces a rapid and strong humoral immune response in ASFV infection and maintains high levels in the blood, saliva, and target organs throughout the infection process[13]. Given its strong antigenicity and immunogenicity, it is a major antigen in the blood of infected pigs[14]. Furthermore, its relatively conserved amino acid sequence makes it a common target for ASFV detection[15]. Encoded by the ASFV E183L gene, P54 is an inner membrane protein involved in the early stages of viral adsorption, invasion, and internalization within hosts[16]. Due to its strong immunogenicity, this protein is widely used in ASFV diagnostic techniques and vaccine development[8,17]. These four proteins were selected for this study based on their critical roles in viral structure, strong immunogenicity, high antigenicity, and significant potential for vaccine development and diagnostic applications.

Because of the threat ASF poses to the global pig industry, the development of ASFV vaccines and diagnostics is necessary, with the identification of B-cell epitopes of ASFV being a critical first step in these processes[18]. Significant progress has been made in identifying B-cell epitopes on CD2v, P72, P30, and P54 proteins. On CD2v, more than 10 distinct B-cell epitopes have been identified, with most of them located in its β-folds[5,19–24]. For P72, which accounts for 31%–33% of the viral mass and forms the ASFV icosahedral capsid, nearly 90 B-cell epitopes have been identified. Some of these epitopes have been found to overlap across various studies[10,25–27]. P72 is stable as a trimer within viral particles, and this higher order structure impacts epitope exposure and antibody recognition[9]. As a result, certain linear B-cell epitopes may be differentially accessible in monomeric versus trimeric forms, leading to variations in epitope profiles reported across studies[28,29]. Although trimeric P72 better reflects native antigenicity, analysis of linear B-cell epitopes using the monomeric form remains important for diagnostic development and subsequent structural studies.[30].P30 and P54 are ASFV inner membrane proteins, which have strong immunogenicity[11]. Over 20 B-cell epitopes on P30 and more than 40 on P54[31–38] have been identified[39–44].

B-cell epitope identification is a crucial step in the creation of disease diagnosis, vaccines, and antibodies[45]. Currently, carrier-based epitope presentation strategies can be broadly divided into two parts. They are fusion protein-based techniques and chemical conjugation methods[46]. Both strategies aim to present epitope peptides on carrier proteins to improve their stability, solubility, and antigenicity[47]. The fusion protein method uses genetic engineering to link epitope peptides to soluble partner proteins like glutathione S-transferase (GST) or thioredoxin (Trx)[48,49]. Based on these tags, the peptides’ production, solubility, and purification are facilitated[50]. In addition, fusion tags enable standardized detection and handling, which is beneficial for high-throughput screening[51]. Chemical conjugation method links synthetic peptide fragments to carrier proteins, such as bovine serum albumin (BSA) or keyhole limpet hemocyanin (KLH), in vitro using chemical cross-linking agents[52]. Gene cloning is not involved in this process, so it is a very adaptable and quick way, especially for peptides that are difficult to express[53]. A variety of coupling chemistries have been developed, including site-specific approaches such as cysteine-maleimide, click chemistry, and iodoacetyl chemistry. These methods allow relatively controlled conjugation through the formation of stable thioether bonds or triazole rings[54,55]. The non-specific methods, like EDC/Sulfo-NHS and the two-step glutaraldehyde crosslinking method, offer broad applicability but may modify multiple sites, potentially affecting epitope orientation. Additionally, oxime/hydrazone ligation has been used for reversible coupling, and azo coupling chemistry has historically been applied for tyrosine-selective labeling[56]. Together, these techniques enable the generation of immunogens or immobilized peptides suitable for applications such as overlapping peptide assays and microarray-based platforms.

However, the current identification of ASFV B-cell epitopes primarily relies on murine immunization protocols. The conventional methodology involves immunizing mice with recombinant ASFV proteins, followed by producing murine-derived monoclonal antibodies (mAbs) via hybridoma technology[10]. Subsequent steps include screening for high-affinity mAbs and identifying epitopes through overlapping peptide mapping[26], with validation ultimately performed using 3 techniques: indirect immunofluorescence assay (IFA), western blotting (WB), and enzyme-linked immunosorbent assay (ELISA)[22,34,57–59]. Notably, only a few of these epitopes (Fig. 1A) have been cross-validated using ASFV-positive porcine sera [22,35,43,60]. B-cell epitopes identified using murine-derived mAbs against specific ASFV proteins may not accurately reflect those targeted by antibodies generated during natural ASFV infection in pigs[61]. Furthermore, conventional detection methods like ELISA and WB rely on hydrophobic interactions that bind protein hydrophobic amino acids to solid surfaces through hydrophobic interactions. This may obscure hydrophobic amino acid residues at the binding interface, compromising assay sensitivity and leading to false-negative results. Shorter peptide epitopes exhibit increasingly stronger masking effects caused by hydrophobic interaction (Fig. 1B)[62]. Therefore, modifying traditional methods like ELISA to overcome these technical limitations is essential for identifying true B-cell epitopes in the context of natural ASFV infection.

**Fig. 1.**
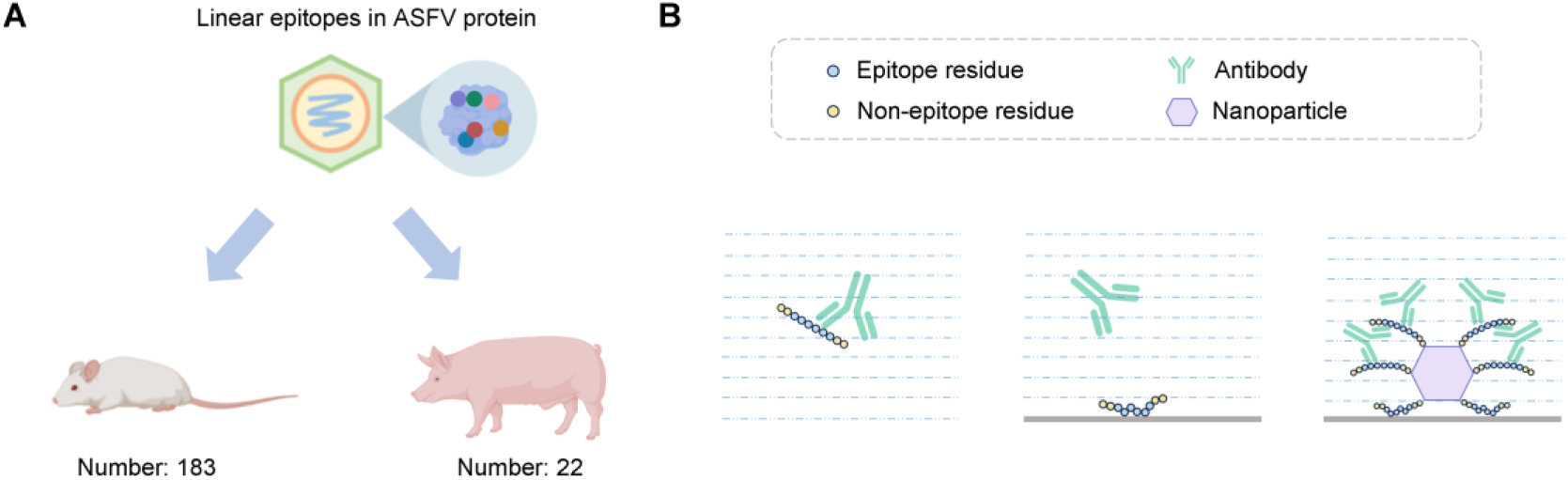
Limitations of conventional ASFV B-cell epitope identification and advantages of ferritin nanoparticle-based epitope identification. **(A)** In the Immune Epitope Database (IEDB), a total of 183 ASFV linear epitopes identified by murine-derived antibodies are registered, among which 22 epitopes have also been validated by porcine ASFV antibodies. **(B)** Comparative schematic of traditional linear epitope identification versus ferritin nanoparticle-based epitope identification. Direct binding of linear epitopes to solid-phase matrices can cause false negatives due to conformational constraints induced by hydrophobic interactions, while ferritin nanoparticles preserve the native conformation and present more epitopes, significantly enhancing detection sensitivity.

Ferritin nanoparticles are iron-storage proteins first identified in 1884. In 1937, Laufberger successfully isolated this protein from horse spleens. This remarkable protein is widespread in biological kingdoms, present in plants, animals, bacteria, and viruses[63]. Ferritin nanoparticles are composed of 20 or 24 subunit monomers, depending on their sizes[64]. Due to their ability to carry drugs internally and be modified with targeting groups externally, ferritin nanoparticles have been widely used in drug delivery, cancer therapy, and bioimaging[65]. The surface of ferritin nanoparticle can stereoscopically display multiple antigenic epitope peptides, which not only avoids site blocking caused by adsorption to solid surfaces but also binds multiple antibody molecules simultaneously, thereby amplifying the binding signal and enhancing the detection sensitivity[66]. However, ferritins from different sources have varying physical and chemical properties. Mammalian ferritins, in particular, may suffer from subunit dissociation and structural instability under certain physiological conditions, which could limit their long-term effectiveness as nanocarriers[67]. However, ferritin from *Pyrococcus furiosus* demonstrates exceptional thermal stability and structural rigidity, with strong subunit interactions[68]. This allows it to remain highly stable under 37°C, effectively preventing the dissociation of its subunits and the breakdown of its nanostructure[69]. Moreover, since *Pyrococcus furiosus* is primarily found in submarine volcanoes, farmed pigs have no specific antibodies against this protein[70]. Because of these properties, we employed *Pyrococcus furiosus* ferritin nanoparticles to display ASFV B-cell epitopes. The high detection sensitivity of recombinant epitope ferritin nanoparticles (Fig. 1B) enables the direct identification of authentic ASFV linear B-cell epitopes using porcine sera.

In this study, we first verified that empty *Pyrococcus furiosus* ferritin nanoparticles do not react with porcine sera. Subsequently, using 7 ASFV P30 B-cell epitopes cataloged in the Immune Epitope Database (IEDB, https://www.iedb.org/) as models, we established an indirect enzyme-linked immunosorbent assay (iELISA) method based on epitope-ferritin nanoparticles, referred to as EFNs-iELISA for short, and confirmed its advantage in detection sensitivity. Employing EFNs-iELISA, we identified multiple B-cell epitopes across 4 key ASFV structural proteins (CD2v, P30, P54, and P72) using ASFV-positive and negative porcine sera. This method enables the identification of immunodominant epitopes relevant to natural ASFV infection, providing crucial insights for the development of next-generation ASFV diagnostics and vaccines.

## 2. Materials and methods

### 2.1 ASFV-positive sera and negative sera

A total of 31 ASFV-negative and 30 ASFV-positive serum samples are from Large White pigs (8 to 10 weeks old) with experimentally infected HLJ/18 ASFV strain (virulent Genotype II). These positive sera were collected at 10 days post-infection (dpi), and then underwent heat inactivation (56℃ for 70 min) in a biosafety level 3 (P3) laboratory at a Company, following standard biosafety protocols. The ASFV-negative sera are from specific pathogen free (SPF) pigs. Besides, all positive and negative serum samples were validated by a commercial blocking ELISA kit (Mingrida, China).

### 2.2 Establishment of the EFNs-iELISA method and validation of ASFV B-cell epitopes

#### 2.2.1 Selection of linear B-cell epitopes from P30

To validate the EFNs-iELISA method, we selected 7 short linear B-cell epitopes (< 20 amino acids) from ASFV P30 for testing. These epitopes (epitope 1 and epitopes 3–8 in Table 1) have all been previously documented in the IEDB.

**Table 1.**
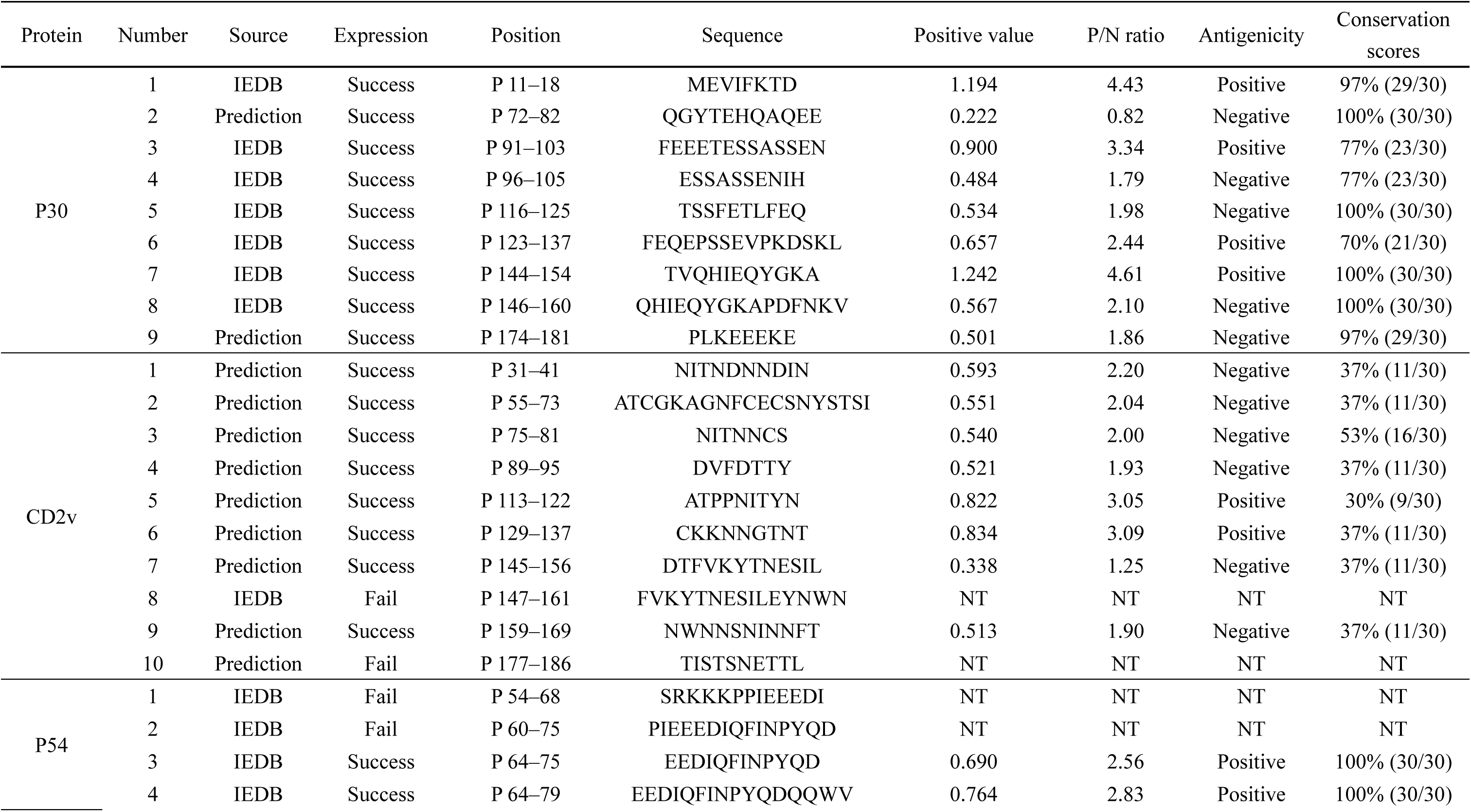
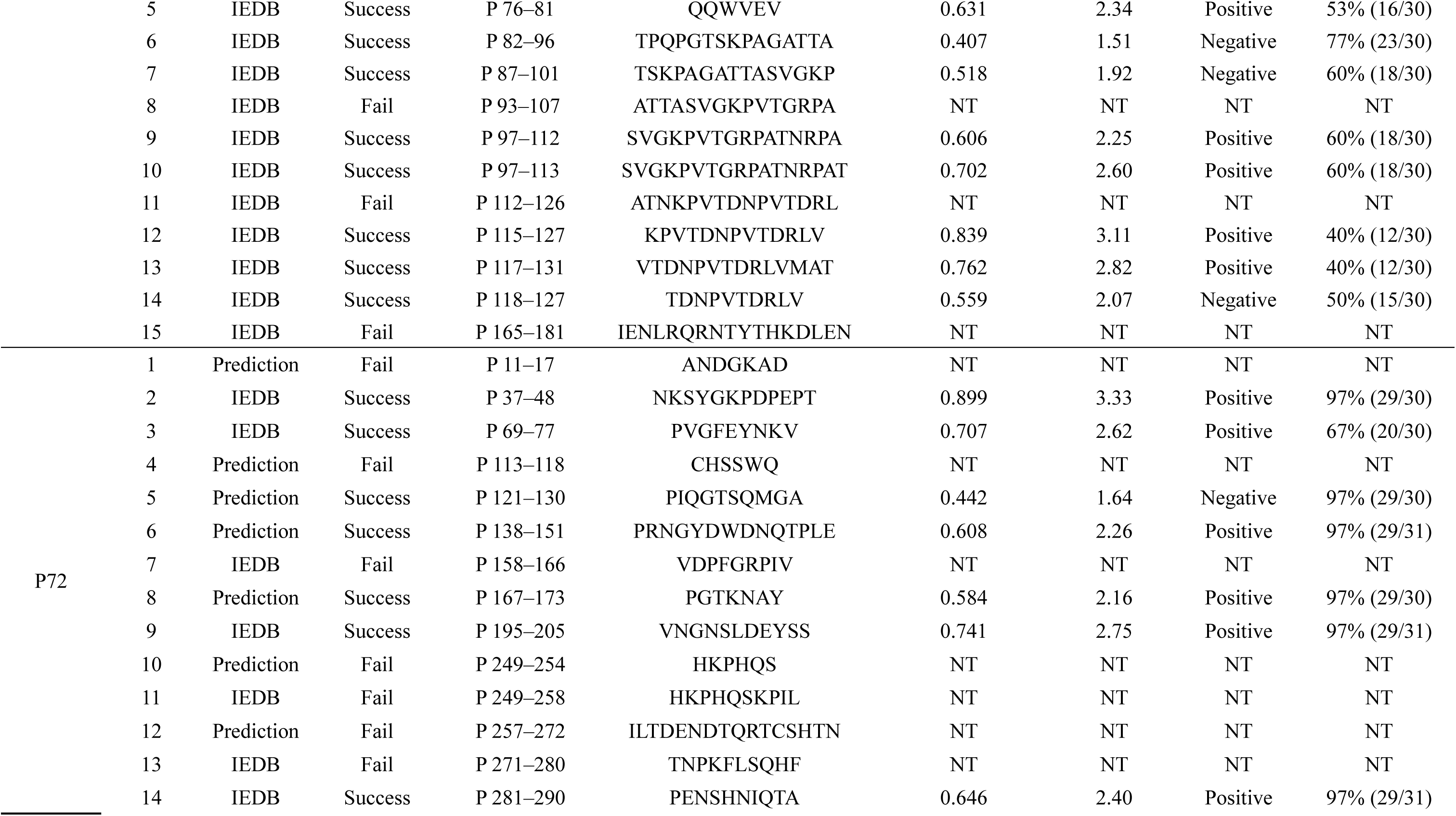
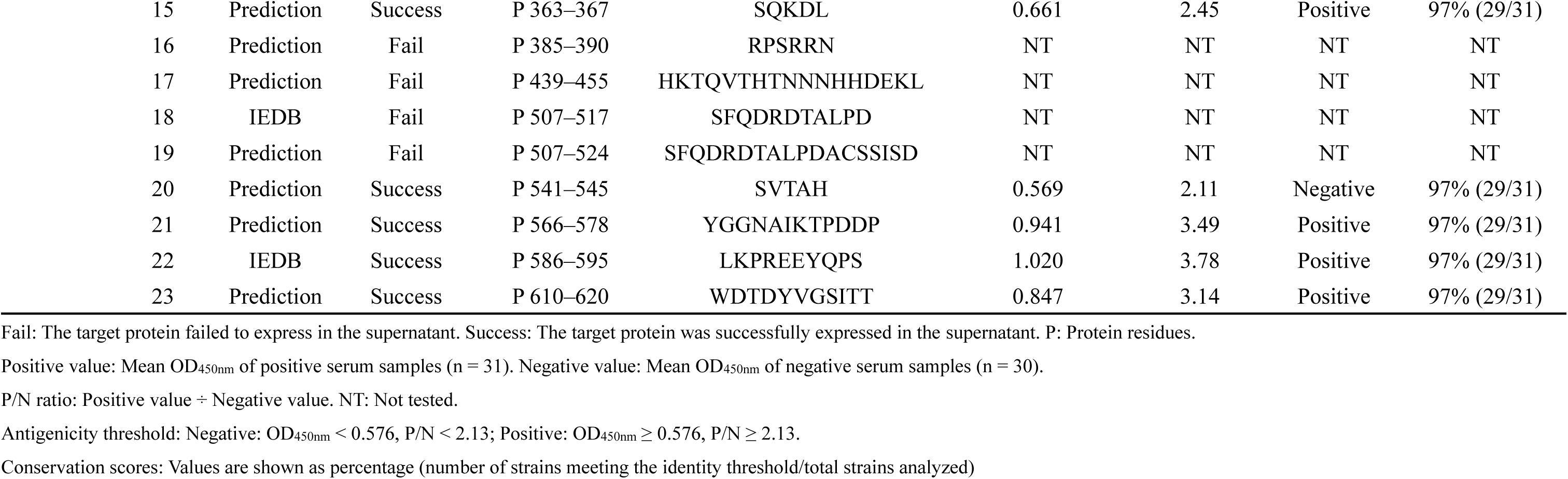
Sequence characterization, expression analysis, and EFNs-iELISA validation of B-cell epitopes from ASFV 4 Proteins.

#### 2.2.2 Expression and purification of EFNs

The ASFV B-cell epitopes were fused to the N-terminal of *Pyrococcus furiosus* ferritin (residues 5–174; GenBank accession no. WP_011011871.1, excluding the signal peptide) via a 2×G_4_S flexible linker. Using a homologous recombination kit (Vazyme, China), the constructs were inserted into the pET-30a(+) vector with XhoI and NdeI restriction enzyme cutting sites, respectively. The recombinant vectors were transformed into BL21(DE3) competent *E. coli* cells (Tsingke, China). Following sequence verification and small-scale induction testing, 200 µL of the bacterial cultures were inoculated into 100 mL of LB medium supplemented with 50 µg/mL kanamycin (Kan+) and grown at 37℃, 200 rpm, shaking for 8 hours. The cultures were scaled up by transferring them to 1.2 L of fresh Kan+ LB medium and continued incubation at 37℃, 160 rpm for 2.5 hours. Protein expressions were induced by adding 0.75 mM IPTG (Solarbio, China), followed by additional incubation at 37℃,160 rpm for 8 hours. The bacterial cells were harvested and lysed by sonication. After heat treatment (70℃, 15 min) and centrifugation, the supernatants containing crude EFNs were collected. The protein solutions were concentrated to 3 mL using a 30 kDa ultrafiltration tube (Millipore, USA). Purification was then performed sequentially on an ÄKTA system (Cytiva, USA). Size-exclusion chromatography was carried out using a Superdex 200 pg 16/60 column (Cytiva, USA) equilibrated with 20 mM Tris-HCl, 50 mM NaCl, pH 8.0, at a flow rate of 1.5 mL/min. This was followed by anion-exchange chromatography on a Resource Q column (Cytiva, USA). The column was equilibrated with buffer A (10 mM Tris-HCl, 10 mM NaCl, pH 8.0). Proteins were eluted with a linear gradient from 0 to 100% buffer B (10 mM Tris-HCl, 1 M NaCl, pH 8.0) over 20 column volumes at a flow rate of 1.0 mL/min. Finally, the purified EFNs were confirmed by SDS-PAGE and transmission electron microscopy before storing them at 4℃.

#### 2.2.3 Blank ferritin antigenicity detection

The 96-well microplates (Thermo, USA) were coated with 100 µL/well of non-epitope-bound ferritin nanoparticles (80, 40, and 20 ng/well) diluted in 50 mM carbonate-bicarbonate buffer (CBS, pH 9.6). After overnight incubation at 4℃, the plates were washed 3 times with PBST (PBS containing 0.05% Tween-20, pH 7.4) using a plate washer at 500 rpm for 3 min per wash. The plates were blocked with 2% BSA-V (Solarbio, China) for 2 hours at 37℃. Next, 31 ASFV-negative and 30 ASFV-positive serum samples were added at 3 dilutions (1:50, 1:100, and 1:200). Different dilutions with different antigen concentrations were combined in pairs; each combination was repeated in 3 independent assays. After incubating at 37℃ for 30 min, the plates were washed, and 100 µL/well of 1:10,000 dilution of rabbit anti-pig IgG-HRP (Bioss, USA) was added. Following another 37℃ incubation for 30 min, the final washes were performed before detection. 100 µL of tetramethylbenzidine substrate (TMB, Biodragon, China) was added to each well and the plate was incubated at room temperature for 15 min. After stopping the reaction with 100 µL TMB stop solution (Biodragon, China), absorbances at 450 nm (OD_450nm_) were measured using a Spark 10 M microplate reader (Tecan, Switzerland). Whether blank ferritin can bind with antibodies in porcine sera specifically was evaluated by comparing OD_450nm_ values and P/N ratios (test sample OD_450nm_ /negative control OD_450nm_) between ASFV-positive and negative sera.

#### 2.2.4 Determination of the optimal antigen coating concentration and serum dilution

Seven constructed P30 EFNs were employed as coating antigens in a checkerboard titration assay to determine the optimal antigen coating concentration and serum dilution. All reagents, instrumentation, and experimental conditions matched those described in section “2.2.3”.

#### 2.2.5 Determination of EFNs-iELISA cut-off threshold

To establish a cut-off threshold for EFNs-ELISA, OD_450nm_ values from 31 ASFV negative serum samples across 39 EFNs (including 7 EFNs described in section “2.2.2” and 32 introduced in section “2.3”) were pooled (n = 1209). The raw OD_450nm_ values were first assessed for normality using D’Agostino & Pearson tests. The results indicated a slight deviation from normality, so a natural log-transformation [ln(OD_450nm_)] was applied to approximate a normal distribution. In the log-transformed data, the mean (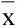) and standard deviation (SD) of negative sera were calculated. The positivity threshold was then calculated as: ln(OD_450nm_) cut-off = 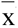 + 2SD. This cut-off value was then back-transformed to the original OD_450nm_ value using the exponential function. Epitopes with average positive OD_450nm_ value ≥ OD_450nm_ cut-off were classified as positive, and those below this value as negative. For P/N ratios’ cut-off classification, the mean OD_450nm_ values of all negative sera(N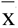) were used as the denominator. The P/N cut-off was calculated as: P/N cut-off = OD_450nm_ cut-off/N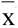. Epitopes with P/N ≥ P/N cut-off were classified as positive, and those with P/N < P/N cut-off were negative.

#### 2.2.6 Epitope validation

Seven P30 EFNs were evaluated by iELISA using optimized antigen coating concentrations and serum dilution with 31 ASFV-negative and 30 ASFV-positive serum samples. Based on OD_450nm_ values and P/N ratios, we confirmed the method’s feasibility for identifying ASFV linear B-cell epitopes.

#### 2.2.7 Comparison with the synthetic peptide iELISA identification method

The P30 P_11–18_, P_96–105_, and P_144–154_ peptides were synthesized with ≥ 95% purity (GenScript, USA). Using these peptides as antigens, iELISA was performed under identical conditions in section “2.2.4”. The OD_450nm_ values and P/N ratios between synthetic peptides and EFNs were compared to show their detection sensitivity.

#### 2.2.8 Construction and expression of GST-His_6_-epitope fusion proteins

Three P30 B-cell epitopes mentioned in section “2.2.7” were fused to the C-terminal of the GST-tag via a 2×G_4_S flexible linker, and there was also a His_6_-tag between GST-tag and linker. The expression procedures followed the same protocol as described in section “2.2.2”. After that, proteins were purified with Ni-NTA agarose resin (Qiagen, Germany) on an ÄKTA system. The column was balanced with binding buffer (20 mM sodium phosphate, 500 mM NaCl, 20 mM imidazole, pH 7.4) and eluted with buffer containing 500 mM imidazole. At last, fusion proteins were coated on the microplates as antigens for iELISA to compare with EFNs-iELISA.

#### 2.2.9 Conjugation of linear epitopes to BSA via EDC/sulfo-NHS coupling

Three epitopes mentioned in section “2.2.7” were conjugated to BSA through EDC/sulfo-NHS coupling, generating traditional fusion protein antigens for iELISA. Firstly, 300 nmol peptides were dissolved in 150 µL 0.1 M MES buffer (pH 5.5–6.0). Then, 10 mM EDC and 25 mM sulfo-NHS were added to activate the peptide carboxyl groups at room temperature, gently stirring for 15 min. The activated peptide solutions were then transferred to BSA (1 mg/mL in PBS, pH 7.4) and incubated overnight at 4 ℃ to form stable amide bonds. At last, the reactions were stopped by adding 20 mM Tris-HCl for 10 min at room temperature. Following conjugation, unreacted small-molecule reagents, including EDC, sulfo-NHS, and MES buffer components, were removed by using a Superdex 200 pg 10/300 column (Cytiva, USA). At last, the BSA-peptide conjugations were coated on the microplates as antigens for iELISA to compare with EFNs-iELISA.

#### 2.2.10 Validation of known murine-derived B-cell epitopes in CD2v, P54, and P72

The known murine-derived linear B-cell epitopes (< 20 amino acids) from ASFV structural proteins (CD2v, P54, and P72) were selected and documented in IEDB (Table 1). Using EFNs-iELISA, these epitopes were validated with ASFV-positive and negative porcine sera.

### 2.3 Prediction and serological screening of immunodominant regions in ASFV CD2v, P30, P54, and P72

The linear B-cell epitopes of ASFV CD2v, P54, and P72 were predicted using 6 different IEDB tools: Bepipred Linear Epitope Prediction 2.0, Chou & Fasman Beta-Turn Prediction, Emini Surface Accessibility Prediction, Karplus & Schulz Flexibility Prediction, Kolaskar & Tongaonkar Antigenicity, Parker Hydrophilicity Prediction (http://tools.iedb.org/main/bcell/). These prediction results were compared with experimentally identified epitopes from ASFV P30, P54, and P72 proteins by EFNs-iELISA (sections “2.2.6” and “2.2.10”). For each prediction method, we respectively calculated the percentages of overlapping amino acid residues within predicted and confirmed epitopes, and then multiplied them to derive the accuracy score for each method’s performance in predicting linear B-cell epitopes (Fig. 5A). Using the prediction tool with the highest accuracy score, we predicted linear B-cell epitopes (5–20 amino acids[71,72]) of ASFV P30, P54, and P72. These epitopes were then identified via EFNs-iELISA (Table 1).

### 2.4 Epitope conservation analysis

For each protein, 30 representative ASFV strains were selected to perform the conservation assessment (Supplementary Table 3 to 6). All epitopes’ conservation was assessed by the “epitope conservancy analysis” tool in IEDB (https://tools.iedb.org/conservancy/). The minimum sequence identity threshold was defined as ≥ 90%. The conservation level of each epitope was expressed as the percentage of strains meeting or exceeding this threshold.

### 2.5 Data analysis

All statistical analyses were performed and graphs were generated using GraphPad Prism 10 (GraphPad Software Inc., CA, USA). For group comparisons, one-way ANOVA was applied with data presented as mean ± standard deviation (SD). The differences statistically significant were considered at *p* < 0.05.

## 3. Results

### 3.1 Sera validation

All serum samples’ blocking ELISA results are shown in Supplementary Table 2.

### 3.2 Determination of EFNs-iELISA cut-off threshold

The ln(OD_450nm_) values from 31 ASFV-negative sera (n = 1209) showed an approximately normal distribution according to the D’ Agostino-Pearson test (*p* > 0.05). The normal Q-Q plot further supported this, showing close alignment between the sample quantiles and the theoretical normal quantiles (Fig. 2A). The ln(OD_450nm_) values also displayed a roughly symmetric, unimodal shape in the frequency histogram (Fig. 2A). Therefore, the cut-off threshold was defined as 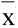 + 2SD in the log-transformed space, yielding ln(OD_450nm_) cut-off = −0.552. Back-transformation resulted in an OD_450nm_ cut-off of 0.576, and the corresponding P/N cut-off was 2.13.

**Fig. 2.**
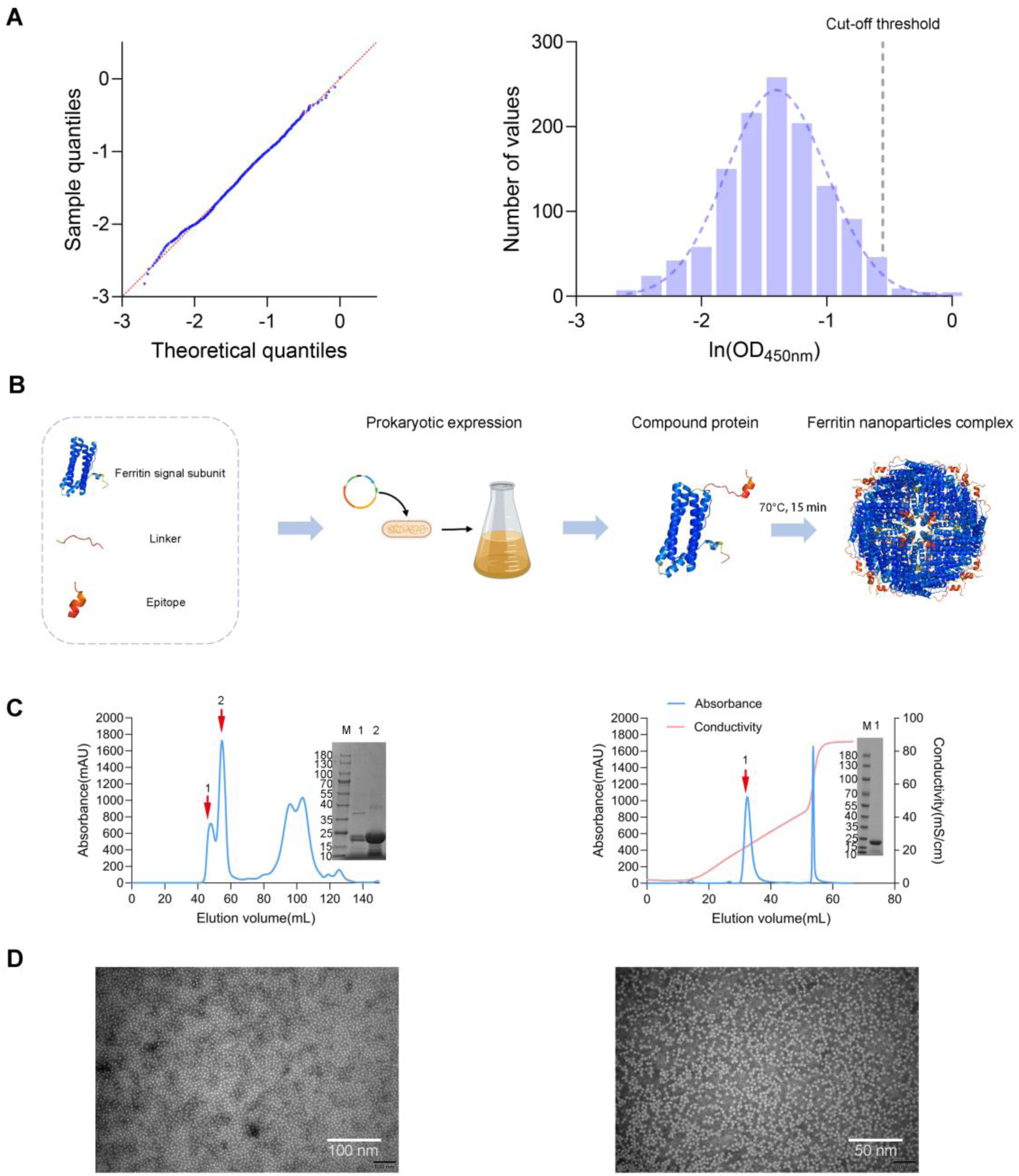
Cut-off threshold calculation, and expression and purification of P30 EFNs. **(A)** Normality assessment of ln(OD₄₅₀) values. Q-Q plot comparing sample quantiles against theoretical normal quantiles (red dashed line). Histogram of ln(OD₄₅₀) with overlaid fitted normal curve (blue dashed line). **(B)** Workflow for EFNs expression. Epitopes were conjugated to *Pyrococcus furiosus* ferritin (monomer ∼19 kDa). **(C)** Purification of P30 P_11–18_–EFNs. Left: size-exclusion chromatography profile with major peaks (red arrows) corresponding to 24-mer nanoparticle complexes (∼450 kDa). Inset: SDS-PAGE analysis confirming monomeric ferritin (∼19 kDa). Right: anion exchange chromatography profile with major peaks (red arrows) corresponding to 24-mer nanoparticle complexes (∼450 kDa). Inset: SDS-PAGE analysis confirming monomeric ferritin (∼19 kDa). **(D)** Transmission electron microscopy (TEM) of P30 P _11–18_–EFNs. Scale bar: 50 nm and 100nm. The inset shows representative 2D class averages demonstrating uniform nanoparticle morphology (24-mer assembly).

### 3.3 Construction of EFNs based on seven known murine-derived P30 B-cell epitopes

The EFNs-iELISA method was established by first constructing a recombinant plasmid encoding a fusion protein in which the B-cell epitope was linked to *Pyrococcus furiosus* ferritin via a flexible linker (Supplementary Fig. 1A). Following expression in a prokaryotic system, the soluble fusion protein was heated to drive 24-subunit ferritin nanoparticles, with the epitopes displayed on the outer surface (Fig. 2B). P30 protein served as a key antigenic target in ASFV infections, and IEDB contains several murine-derived linear B-cell epitopes for P30 (Table 1). Considering the typical 5-15 amino acid length range for B-cell epitopes[73], we chose 7 epitopes (< 20 amino acids) for EFNs construction. Using EFNs-iELISA, we assessed whether murine-derived B-cell epitopes could specifically bind antibodies in porcine sera. This not only validated the EFNs-iELISA’s functionality but also evaluated the reality of murine-derived epitopes.

Seven P30 EFNs were successfully constructed. As illustrated by the P_11–18_–EFNs representative (Fig. 2B), the EFNs were expressed efficiently in E. coli and remained soluble. After heat treatment, size-exclusion chromatography demonstrated a predominant high-molecular-weight peak, which was subsequently purified into a single peak through anion-exchange chromatography. These results confirmed that the recombinant proteins had self-assembled the EFNs through heating (Fig. 2C). To further confirm nanoparticle formation, we characterized the purified protein using transmission electron microscopy (TEM). The TEM images revealed well-dispersed, spherical EFNs in solution (Fig. 2D). These results demonstrated successful EFNs assembly and purification. The expression and purification profiles for the remaining 6 P30 EFNs are shown in Supplementary Fig. 2A.

### 3.4 Validation of the EFNs-iELISA method and ASFV P30 B-cell epitopes

The absence of specific reactivity between *Pyrococcus furiosus* ferritin and porcine sera was essential for reliable ASFV B-cell epitope identification. We first evaluated blank ferritin nanoparticles using iELISA with 30 ASFV-positive and 31 negative serum samples (Fig. 3A). Both positive and negative groups showed similarly low OD_450nm_ values (*p* > 0.05) with P/N ratios < 1.5 (Fig. 3B, Supplementary Table 1), demonstrating no specific antibody to blank ferritin in swine populations. This confirmed the ferritin nanoparticle’s suitability as an epitope display platform without interfering with ASFV-specific antibody detection.

**Fig. 3.**
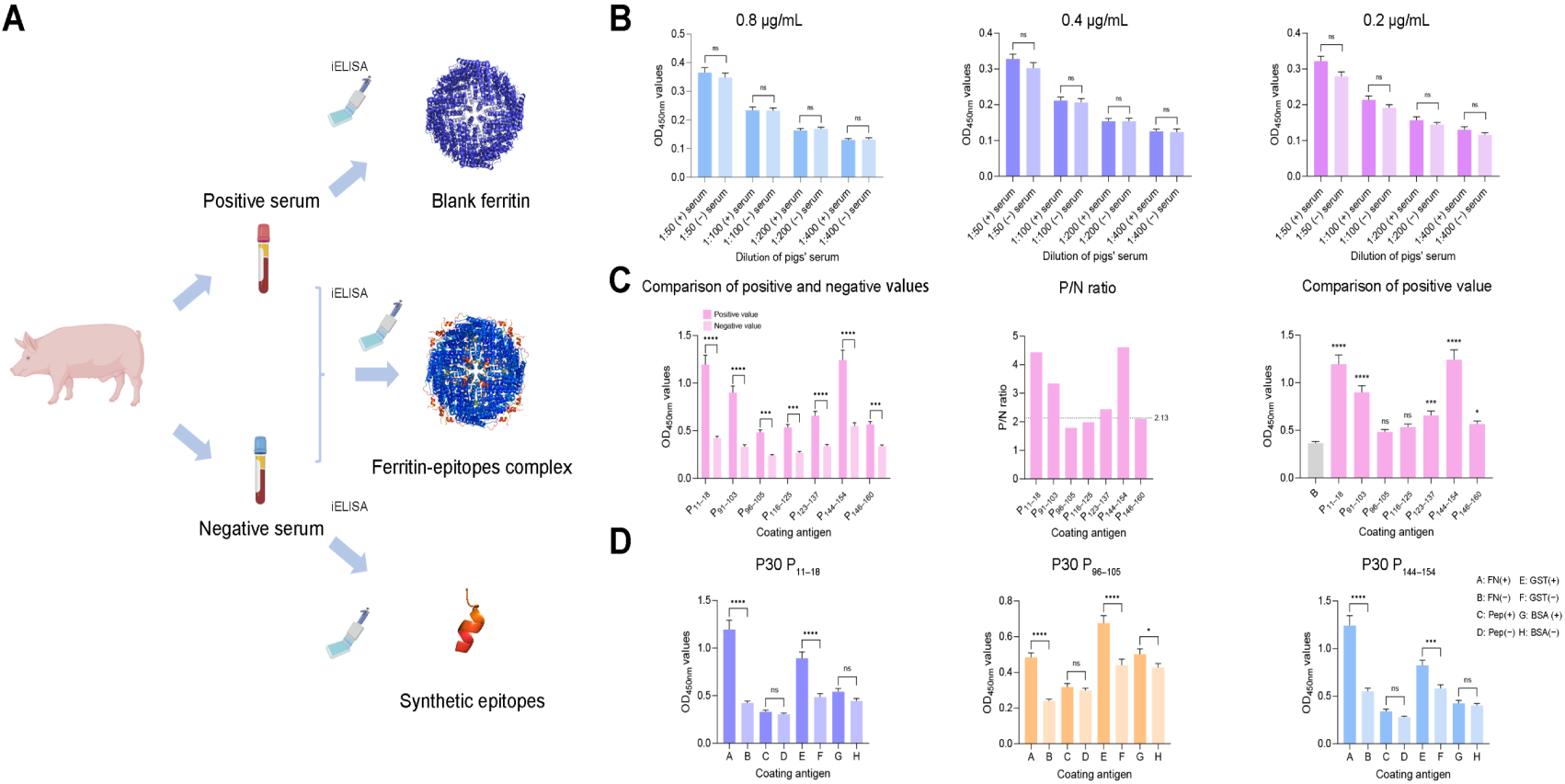
Validation of the EFNs-iELISA method. **(A)** Process of validation of the EFNs-iELISA method. **(B)** Blank ferritin antigenicity detection with the EFNs-iELISA method. Data = mean ± SD; Positive sera n = 30; negative sera n = 31; ns: *p* > 0.05 (ANOVA). **(C)** Validation of P30 B-cell epitopes with the EFNs-iELISA method. Data = mean ± SD; Positive sera n = 30; negative sera n = 31; ns: *p* > 0.05; *: *p* < 0.05; ***: *p* < 0.001; ****: *p* < 0.0001 (ANOVA). (D) Comparative performance between the traditional epitope iELISA detection methods and EFNs-iELISA method in P30 epitope identification. Data = mean ± SD; Positive sera n = 30; negative sera n = 31; ns: *p* > 0.05; ****: *p* < 0.0001 (ANOVA). FN: EFNs, Pep: synthetic peptide, GST: GST-His_6_-epitope, BSA: BSA-conjugated epitope.

The iELISA comparison of OD_450nm_ values between ASFV-positive and negative porcine sera against EFNs and blank ferritin enabled immunodominant regions screening (Fig. 3C). Through a checkerboard titration assay, we determined 0.8 µg/mL coating concentration and 1:50 serum dilution as optimal conditions, where EFNs showed maximal P/N ratios with positive OD_450nm_ values approaching 1.000 (Supplementary Fig. 2B). All subsequent iELISA experiments employed the same optimal conditions. Notably, all EFNs demonstrated significantly higher positive OD_450nm_ values compared to negative OD_450nm_ values (*p* < 0.05). However, the OD_450nm_ values varied across different EFNs. For example, P_11–18_–EFNs and P_144–154_–EFNs exhibited particularly strong signals with OD_450nm_ values exceeding 1.0 in positive sera, while other EFNs’ positive OD_450nm_ values are smaller than 1.0. Based on P/N ratios, P_11–18_–EFNs, P_91–103_–EFNs, P_123–137_–EFNs, and P_144–154_–EFNs demonstrated positive reactivity (P/N ≥ 2.13), while others showed negative reactivity (P/N < 2.13) (Table 1). The positive OD_450nm_ values of P30 EFNs compared with blank ferritin nanoparticles showed highly significant differences for P_11–18_–EFNs, P_91–103_–EFNs, P_123–137_–EFNs, and P_144–154_–EFNs, while no significant differences were observed for P_96–105_–EFNs and P_116–125_–EFNs (Fig. 3C). Based on the above analysis, we considered P_11–18_, P_91–103_, P_123–137_, and P_144–154_ of the P30 protein as positive B-cell epitopes, while the remaining epitopes are negative.

### 3.5 Comparative analysis of EFNs-iELISA with traditional epitope iELISA detection methods

Proteins were expressed and constructed successfully, and the purification results were shown in Supplementary Fig. 2D. As shown in Fig. 1B, we believed EFNs-based identification of ASFV immunodominant B-cell regions reduced false positives caused by spatial steric hindrance compared to synthetic peptides, GST-His_6_-peptides and BSA-conjugated peptides, while antibody enrichment enhances detection sensitivity. To validate this, we synthesized and constructed positive epitopes (P_11–18_, P_144–154_) and a negative epitope (P_96–105_) in three different formats, performed iELISA, and compared the results with EFNs-iELISA identification (Fig. 3D).

The results showed no significant difference in OD_450nm_ values between positive and negative groups for all synthetic peptides and BSA-conjugated P_11–18_, P_144–154_ (*p* > 0.05). In contrast, both EFNs and GST-His_6_-peptides, exhibited significantly higher positive OD_450nm_ values than negative OD_450nm_ values (*p* < 0.0001 and *p* < 0.001). Furthermore, EFNs showed significantly higher positive OD_450nm_ values than the synthetic peptides, GST-His_6_-peptides and BSA-conjugated peptides’ OD_450nm_ values. The only exception was in the P_96–105_ group, where GST-His_6_-P_96–105_ and BSA-conjugated P_96–105_ had higher positive OD_450nm_ values than EFNs’ positive values, indicating a high background signal.

This structural advantage translated into markedly improved assay performance, yielding a 2-fold higher signal-to-noise ratio than the synthetic-peptide iELISA and approximately a 1.2-fold improvement over the GST-HIS_6_-peptide and BSA-conjugated peptide assays. GST-His_6_-epitopes and BSA-conjugated-epitopes had lower discriminative ability on some occasions because of the high background signal. These findings demonstrated that EFNs-iELISA exhibits superior sensitivity compared to traditional iELISA, while effectively preventing false-negative results and enhancing detection accuracy.

### 3.6 Validation of known murine-derived B-cell epitopes in CD2v, P54, and P72

Beyond P30, CD2v, P54, and P72 are also important antigenic proteins in ASFV, and they may induce the production of neutralizing antibodies[74]. Therefore, we next applied our newly developed method to validate linear B-cell epitopes in these proteins. From the 3 proteins’ linear B-cell epitopes in the IEBD, we selected epitopes shorter than 20 amino acids (Table 1) and constructed their EFNs. While most EFNs were successfully generated (Supplementary Fig. 3), some EFNs failed to express, including the sole CD2v epitope (Table 1).

EFNs-iELISA results (Fig. 4A) revealed significantly higher EFNs’ positive OD_450nm_ values than negative OD_450nm_ values and also than blank ferritin nanoparticles tested with ASFV-positive sera. All EFNs of P72 displayed positive reactivity, with a P/N ratio ≥ 2.13. This result indicates that all IEDB-selected linear B-cell epitopes of P72 are positive.

**Fig. 4.**
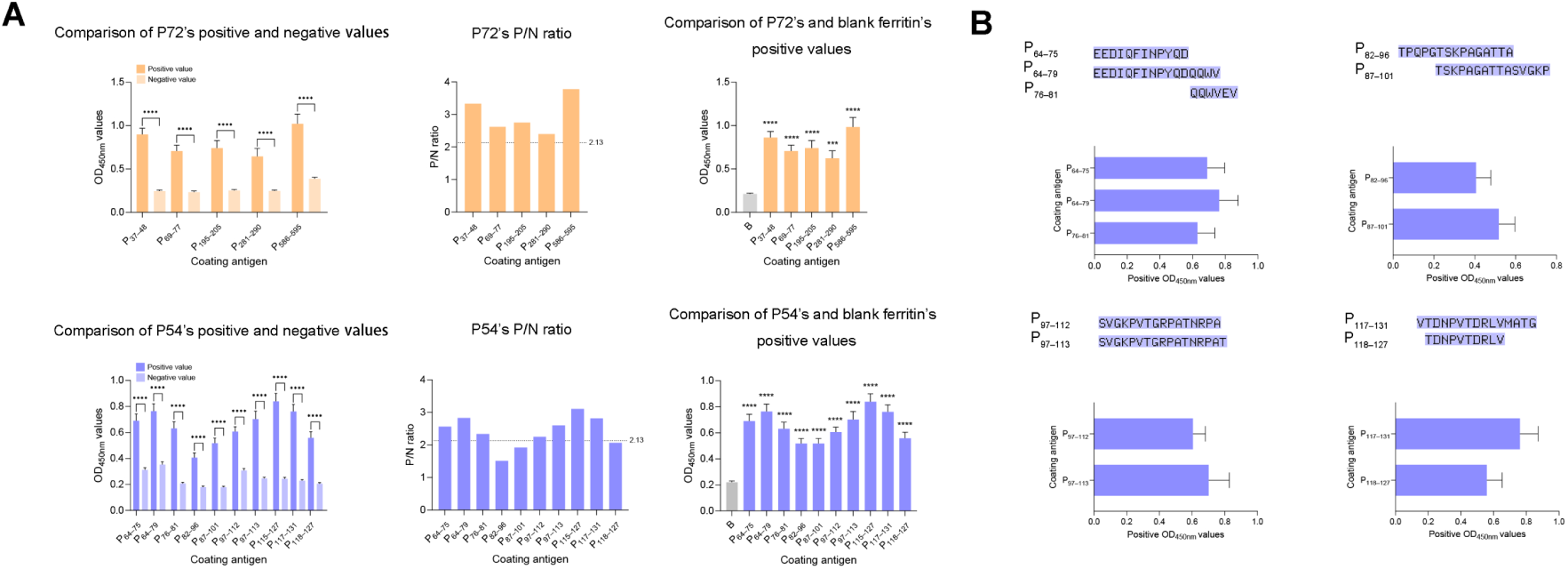
Validation of known murine-derived B-cell epitopes in CD2v, P54, and P72 with the EFNs-iELISA method. **(A)** Results of the EFNs-iELISA method. Data = mean ± SD; Positive sera n = 30; negative sera n = 31; ***: *p* < 0.001; ****: *p* < 0.0001 (ANOVA). **(B)** Antigenicity comparison between overlapping epitopes of P54. Data = mean ± SD; Positive sera n = 30.

For P54, the positive OD_450nm_ values of EFNs varied among epitopes (0.407–0.839), reflecting differences in their reactivity. When evaluated by P/N ratios, P_64–75_–EFNs, P_64–79_–EFNs, P_76–81_–EFNs, P_97–112_–EFNs, P_97–113_–EFNs, P_115–127_–EFNs, and P_117–131_–EFNs showed positive reactivity (P/N ≥ 2.13). In comparison, P_82–96_–EFNs, P_87–101_–EFNs, and P_118–127_–EFNs show negative reactivity (P/N < 2.13). Notably, P54 contains multiple overlapping epitopes (P_64–75_/P_64–79_/P_76–81_, P_97–112_/P_97–113_, P_82–96_/P_87–101_, and P_117–131_/P_118–127_). There is a strong correlation between epitope length, their overlapping relationships, and positive OD_450nm_ values, as exemplified by P_64–75_/P_64–79_/P_76–81_, P_97–112_/P_97–113_, and P_117–131_/P_118–127_ (Fig. 4B). This is because longer epitopes are more likely to cover key antigenic regions, maintain better conformational integrity, and thereby exhibit stronger antibody-binding capacity. These findings demonstrated that EFN-iELISA offers both the sensitivity and resolution required for precise epitope identification.

### 3.7 Prediction and serological screening of immunodominant regions in ASFV CD2v, P30, P54, and P72

Only one CD2v epitope was cataloged in the IEDB (Table 1), indicating that numerous epitopes within these key ASFV antigenic proteins remain to be identified. To address this, we tried to predict new linear B-cell epitopes and construct EFNs, and then verified them. Using 6 distinct prediction methods from IEDB (http://tools.iedb.org/main/bcell/), we observed variability among the different methods’ outputs. To identify the most reliable prediction method, we systematically evaluated all 6 algorithms against our 22 experimentally confirmed epitopes. Prediction accuracy was quantified by measuring the concordance between computational predictions and empirically identified epitopes (Fig. 5A). Our comparative analysis revealed that Bepipred Linear Epitope Prediction 2.0 achieved the highest accuracy (17.74%), closely followed by Parker Hydrophilicity Prediction (17.53%) (Fig. 5A). However, Bepipred 2.0 tended to generate excessively long, imprecise epitope predictions (Supplementary Fig. 2C). Based on this trade-off between accuracy and precision, we selected Parker Hydrophilicity Prediction for subsequent analyses. Using prediction analysis, we discovered 25 novel linear B-cell epitopes across P30, P72, and CD2v (Table 1). There were no additional epitopes predicted for P54, as the existing identified epitopes for this protein were already comprehensive.

Following prokaryotic expression and purification, we successfully constructed 17 EFNs (Table 1 and Supplementary Fig. 3B). EFNs-iELISA analysis (Fig. 5B, Table 1) showed that the P30 P_72–82_ and P_174-181_ exhibited consistently negative serological reactivity across multiple evaluation metrics (positive OD_450nm_ values and P/N ratios). Among the 8 predicted CD2v epitopes, P_31–41_–EFNs, P_113–122_–EFNs and P_129–137_–EFNs exhibited positive reactivity (P/N ≥ 2.13). While the other five epitopes exhibited weak antigenicity, with reactivity levels under the negative threshold (P/N < 2.13). Among the peptides classified as non-reactive, P_145–156_–EFNs displayed extremely low positive OD_450nm_ value, and had no significant differences between blank ferritin OD_450nm_ value across positive sera (*p* > 0.05). For P72’s predicted epitopes, only P_121–130_–EFNs and P_541–545_–EFNs showed negative serological reactivity (P/N < 2.13). This result showed the same trend with positive OD_450nm_ values in comparison between blank ferritin and these two epitopes. Overall, 8 predicted epitopes across these 3 proteins are immunodominant regions.

### 3.8 Epitope conservation analysis

In P30 epitopes P_11–18_, P_72–82_, P_116–125_, P_144–154_, P_146–160_, and P_174–181_ were confirmed as highly conservation epitopes, defined as being present in ≥ 90% of the analyzed protein sequences. In CD2v, all epitopes showed low conservation, with their conservation rates falling below 60%. In P54, only epitopes P_64–75_ and P_64–79_ showed high conservation (100%). All other epitopes exhibited variable and generally lower conservation, ranging from moderate (60% to 77%) to low levels (40% to 53%). In P72, all epitopes were highly conserved except for P_69–77_, which exhibited a moderate conservation level (67%).

### 3.9 Comparative analysis of ASFV murine-derived B-Cell epitopes and porcine-derived B-Cell epitopes

As shown in the Fig. 6 and Table 1, among the 22 murine-derived positive B-cell epitopes from P30, P54, and P72, 16 were confirmed as immunodominant B-cell regions in naturally ASFV-infected pigs. Approximately 72% of the murine-derived positive epitopes overlapped with porcine-derived positive epitopes, suggesting that despite species differences, the identified murine-derived ASFV B-cell epitopes still have considerable credibility. However, approximately 30% of murine-derived epitopes failed to show strong reactivity with ASFV-positive porcine sera. Furthermore, some murine-derived epitopes in IEBD are excessively long and potentially contain multiple epitopes, showing overlap with the 12 porcine-derived ASFV immunodominant B-cell regions we identified in this study.

**Fig. 5.**
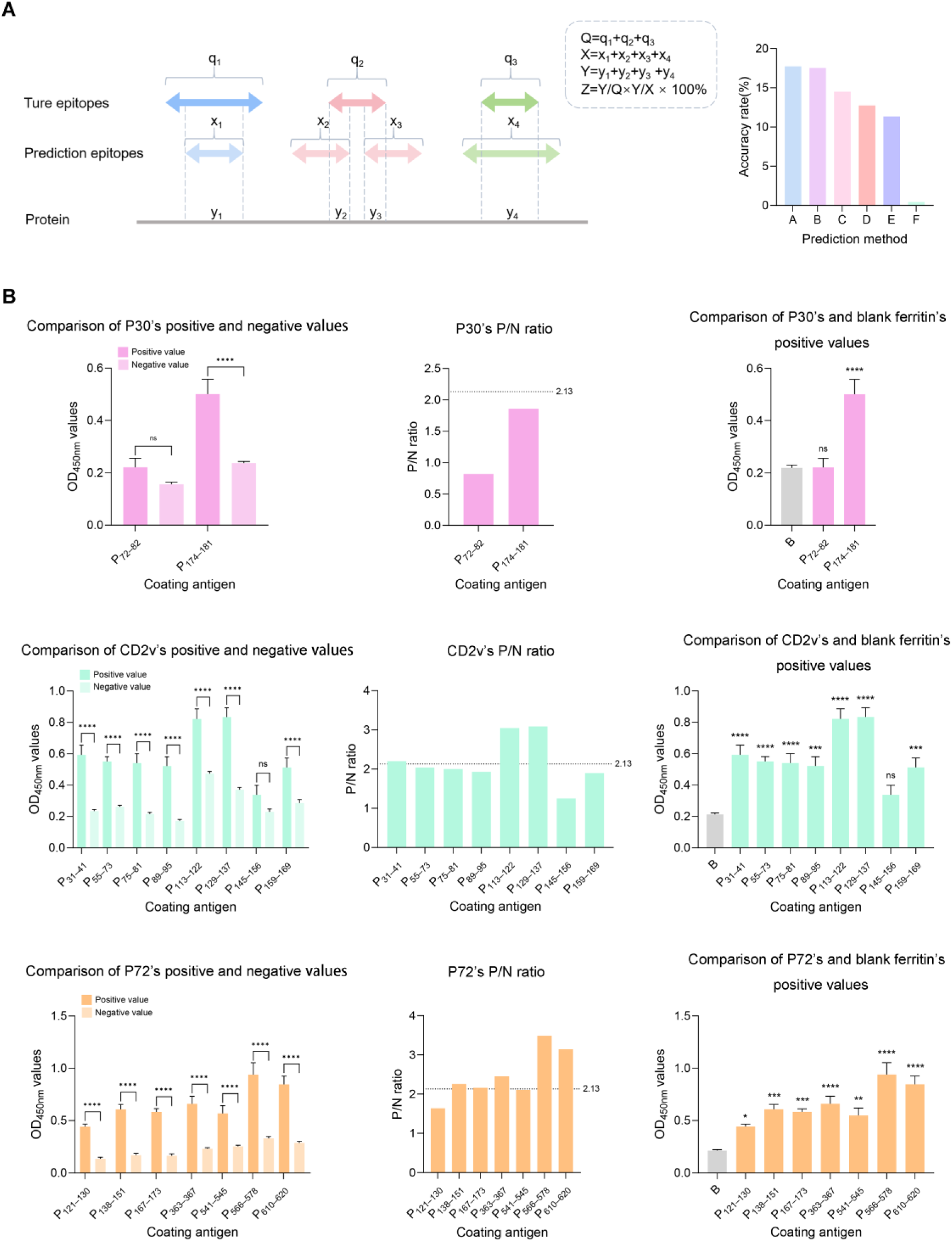
Computational prediction, EFNs-iELISA method screening immunodominant regions in ASFV P30, CD2, and P72. **(A)** Accuracy calculation of each epitope prediction technology. Q: Number of amino acids of identified epitopes in P30, CD2v, and P72. X: Number of amino acids of predicted epitopes in P30, CD2v, and P72. Y: Number of overlapping amino acids between predicted and identified epitopes in P30, CD2v, and P72. Y/Q: The percentage of overlapping amino acids in relation to the total amino acid count of the identified epitope. Y/X: The percentage of overlapping amino acids in relation to the total amino acid count of the predicted epitope. Z: Accuracy rates of each prediction technique when predicting ASFV CD2v, P54, and P72 B-cell epitopes. **(B)** Results of the EFNs-iELISA method. Data = mean ± SD; Positive sera n = 30; negative sera n = 31; ns: *p* > 0.05; *: *p* < 0.05; **: *p* < 0.01; ***: *p* < 0.001; ****: *p* < 0.0001.

**Fig. 6.**
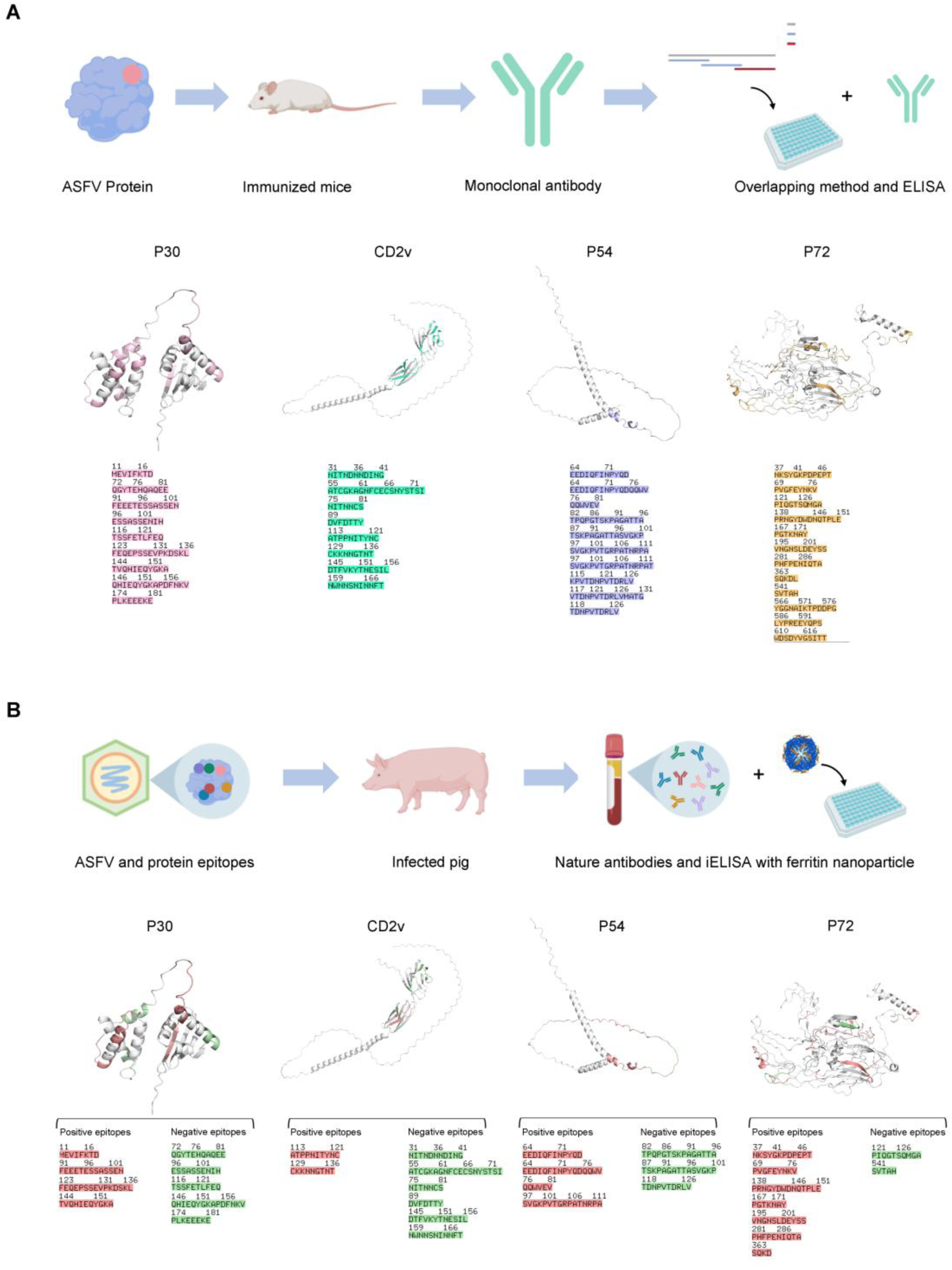
Comparative analysis of B-cell epitope screening strategies and structural epitope visualization. **(A)** Identification of ASFV linear B-cell epitopes by murine-derived mAbs. **(B)** Screening and validation of ASFV linear B-cell epitopes and immunodominant regions by the EFNs-iELISA method. The PDB code for P72 is 612t. Structures of P30, CD2v, and P54 were predicted by AlphaFold3.

By integrating structural data from AlphaFold3 predictions and experimental analyses, we mapped the spatial distribution of these epitopes. Most epitopes were exposed on the surface of the proteins. However, when the P72 molecule forms a trimer, P_566–578_ and P_586–595_ are completely hidden inside. Most epitopes localize to structured protein regions (α-helices or β-sheets), while a few are located on disordered regions (primarily loops and turns), as seen in P_91–101_ of the P30 and P_76–81_, P_97–112_, P_97–113_, P_115–127_, and P_117–131_ in P54. Antibodies generated against these linear epitopes demonstrated reactivity against both native ASFV proteins and intact viral particles, suggesting their potential worth.

## 4. Discussion

ASF, a highly lethal and contagious disease, severely threatens global pig industry economics and biosecurity[1,2]. Currently, no effective vaccines or treatments exist for ASFV. Identification of ASFV B-cell epitopes remains fundamental for developing diagnostic tools and vaccines[18]. CD2v, P30, P54, and P72 exhibit strong antigenicity and immunogenicity [9,11], making screening and serological evaluation of their immunodominant regions valuable for ASFV prevention and control. However, most reported ASFV B-cell epitopes were defined based on murine-derived mAbs, and their reactivity with naturally infected pigs’ antibodies remains incompletely characterized. To address this limitation, immunodominant regions can be screened and validated using ASFV-positive and ASFV-negative porcine sera. The iELISA has been well-established as a high-throughput, cost-effective method for protein antigen screening. It faces limitations in precisely identifying specific epitopes, especially the short peptide epitopes. This challenge arises because serum antibody concentrations and affinities against individual epitopes frequently fall below the detection threshold (Fig. 3D)[61,62]. The ferritin nanoparticles derived from the hyperthermophilic archaeon *Pyrococcus furiosus* exhibit remarkable structural stability and self-assembly properties, along with signal amplification capabilities. These nanoparticles have been extensively utilized in drug delivery systems and vaccine development platforms[65,75]. By incorporating ferritin nanoparticles to address the technical limitations of iELISA in B-cell epitope mapping, this method enables rapid, cost-effective, and precise screening and validation of ASFV immunodominant regions. This advancement will significantly facilitate the development of novel vaccines and diagnostic reagents against ASFV.

During the development of the EFNs-iELISA method, we first confirmed that blank ferritin nanoparticle carriers showed negative reactivity to porcine sera (Fig. 3B, Supplementary Table 1), thereby ensuring the sensitivity and specificity of subsequent epitope identification. Using 7 murine-derived B-cell epitopes from the ASFV P30 as a model, EFNs-iELISA successfully differentiated these epitopes based on their reactivity with ASFV-positive and negative porcine sera, enabling quantitative antigenicity assessment (Fig. 3C).

In the comparison experiments with synthetic peptides, GST-HIS_6_-epitope and BSA-conjugated peptides, EFNs significantly improved B-cell epitope detection sensitivity through three-dimensional, multivalent epitope display (Fig. 3D). The synthetic peptides’ weak antigenicity has two reasons. First, when binding with microplates, synthetic peptides often present flattened orientations that expose non-epitope regions. Second, these three P30 epitopes we selected (8 to 11 amino acids) are too short to present high antigenicity. This reduced specific binding explains the small differences between positive and negative OD_450nm_ values. The GST-HIS_6_-epitopes have larger positive OD_450nm_ values compared to synthetic peptides, because GST-HIS_6_ tags can improve the epitope’s absorption and stabilization. However, as a large heterologous protein, GST can interact with nonspecific IgG and anti-GST antibody, which causes the high background signal. BSA-conjugated epitopes showed unexpectedly weak antigenicity and poor distinguishability between positive and negative epitopes. This result is mainly attributed to limitations of EDC/sulfo-NHS coupling and short peptides. The EDC/sulfo-NHS coupling may activate carboxyl groups on the peptide C-terminus or Asp/Glu side chains, resulting in random conjugation to lysine residues on BSA and masking key epitope residues. The short peptides’ binding efficiency is often low and heterogeneous, leading to low peptide density on BSA molecules, and improving the background signal. Overall, compared to synthetic peptides, GST-HIS_6_-epitope and BSA-conjugated peptides, EFNs can display short epitopes ordered and multivalently. This increased the sensitivity and specificity in linear B-cell epitopes’ screening and made EFNs-iELISA an efficient method.

Using EFNs-iELISA, we validated that 72% murine-derived epitopes also have positive reactivity with ASFV naturally infected sera. We also screened 7 immunodominant regions on ASFV structural proteins CD2v, P30, P54 and P72 (Fig. 6, Table 1). Based on conservation scores and serological reactivity, several epitopes from P30 (P_11–18,_ P_91–103_, P_123–137_, and P_144–154_), P54 (P_64–75_ and P_64–79_), and P72 (P_37–48_, P_195–205_, P_281–290,_ and P_586–595_) were validated as promising candidates for chimeric nanoparticle vaccine development and serological diagnostics. In addition, several immunodominant regions on P72 (P_167–173_, P_138–151_, P_363–367_, P_566–578_, and P_610–620_) were identified through serological reaction as candidates for further investigation. These results revealed that stronger epitope reactivity was associated with localization in β-sheet and flexible loop regions, consistent with increased surface accessibility. Notable differences were observed between the murine-derived and porcine-derived epitope profiles. These discrepancies likely arise from 2 key factors. Firstly, immunization of mice with individual ASFV proteins eliminates interprotein antigenic competition, potentially enhancing epitope detection sensitivity for the immunized antigens. However, during natural ASFV infection, the full complement of viral epitopes competes for immune recognition, resulting in only immunodominant epitopes eliciting strong serological responses. Secondly, during mAb-based epitope identification, the selection process favors mAbs with superior binding affinity, effectively amplifying the detected reactivity of target epitopes. In contrast, reactivity with ASFV-positive porcine sera reflects the average binding affinity across the entire antibody repertoire present in naturally infected serum. Therefore, EFNs-iELISA using porcine serum provides an accurate quantitative assessment of ASFV epitope antigenicity. This approach enables reliable screening and validation of immunodominant regions, significantly advancing the development of precision vaccines and diagnostic reagents against ASFV.

Our study also showed several limitations of the EFNs-iELISA method. First, during EFNs’ construction, some of them failed to express. Such failures may be triggered by structural incompatibility between certain epitopes and the ferritin subunit, instability of the fusion protein in E. coli or poor solubility of compound proteins. This reflects the limitations of EFNs platform, that expression and purification need case-by-case optimization, and the workflow does not support high-throughput screening. We may optimize the EFNs workflow and explore other scaffolds (SpyTag/SpyCatcher nanoparticles, virus-like particles) in further studies. Obtaining highly pure nanoparticles still requires sequential purification through size-exclusion and ion-exchange chromatography to remove excess unassembled subunits. Future methodological improvements, such as implementing simplified ultrafiltration approaches, could streamline both the construction and purification processes, potentially enhancing the technology’s scalability and widespread adoption. EFN-iELISA also has advantages compared with high-throughput epitope detection methods such as phage display and peptide arrays. Phage display can select binding peptides efficiently, but it often identifies murine-derived viral epitopes. Peptide arrays are expensive and limited by specialized instrumentation. Compared with these methods, EFN-iELISA provides a balance of authenticity, cost, and accessibility. These combined benefits significantly lower the technical and economic barriers to implementation, making the method promising for widespread applications.

In summary, we have developed the EFNs-iELISA for the validation of B-cell epitopes and immunodominant regions’ screening on 4 key ASFV structural proteins (CD2v, P30, P54, and P72). This method provides both technical innovation and critical epitope data to advance ASFV vaccine design and diagnostic development. EFNs-iELISA provides a practical and stable approach for serological screening and validation of pathogen-derived B-cell regions, supporting applications in viral immunology research.

## Acknowledgements

This study was supported by National Key Research and Development Program of China (2021YFD1800100), National Natural Science Foundation of China (32172871), 2115 Talent Development Program of China Agricultural University. We would like to thank Dr. Jie Li, Dr. Teng Teng, Xingqi Zou, Dr. Kai Wen and Inner Mongolia Jinyu Company for providing the ASFV-positive and negative porcine sera. During the preparation of this essay, the authors used ChatGPT to check grammar and improve language. After using the tool, we reviewed and edited the content as needed and took full responsibility for the content of the publication.

## Notes

### Competing Interest Statement

The authors have declared no competing interest.

