## Supplementary Table 1 to Table 6, Supplementary Fig. 1 to Fig. 3 for "Ferritin Nanoparticle-Based Indirect ELISA for Immunodominant Region Screening and B-cell Epitope Validation of African Swine Fever Virus"

**Supplementary Material**

**Supplementary Table 1 Blank ferritin nanoparticles’ antigenicity detection.**

Coating concentration

| OD_450nm_ | | 0.8 | 0.4  （μg/mL） | 0.2 |
| --- | --- | --- | --- | --- |
| 1:50  Serum dilution | P | 0.366 | 0.328 | 0.322 |
|  | N | 0.349 | 0.302 | 0.279 |
|  | P/N | 1.0 | 1.1 | 1.2 |
| 1; 100 | P | 0.234 | 0.212 | 0.213 |
|  | N | 0.233 | 0.206 | 0.191 |
|  | P/N | 1.0 | 1.0 | 1.1 |
| 1: 200 | P | 0.163 | 0.154 | 0.157 |
|  | N | 0.169 | 0.154 | 0.144 |
|  | P/N | 1.0 | 1.0 | 1.1 |
| 1: 400 | P | 0.131 | 0.126 | 0.13 |
|  | N | 0.132 | 0.123 | 0.116 |
|  | P/N | 1.0 | 1.0 | 1.1 |

P: Positive value, mean OD_450nm_ of positive serum samples (n = 31).

N: Negative value, mean OD_450nm_ of negative serum samples (n = 30).

P/N ratio: Positive value ÷ Negative value.

**Supplementary Table 2 Serum samples detection results with blocking ELISA kit**

| Positive serum number | Value | Negative serum number | Value |
| --- | --- | --- | --- |
| 1 | 61.64% | 1 | 4.82% |
| 2 | 71.66% | 2 | 16.20% |
| 3 | 83.19% | 3 | 15.05% |
| 4 | 76.43% | 4 | 38.52% |
| 5 | 85.66% | 5 | 8.94% |
| 6 | 84.39% | 6 | 12.54% |
| 7 | 61.70% | 7 | 17.20% |
| 8 | 87.93% | 8 | 0.59% |
| 9 | 90.58% | 9 | 23.33% |
| 10 | 57.97% | 10 | 22.65% |
| 11 | 81.80% | 11 | 4.32% |
| 12 | 61.90% | 12 | 8.90% |
| 13 | 60.70% | 13 | 12.02% |
| 14 | 94.50% | 14 | 5.52% |
| 15 | 76.80% | 15 | 13.90% |
| 16 | 80.41% | 16 | 21.57% |
| 17 | 59.37% | 17 | 16.60% |
| 18 | 66.13% | 18 | 19.49% |
| 19 | 57.28% | 19 | 8.95% |
| 20 | 74.09% | 20 | 10.82% |
| 21 | 68.66% | 21 | 42.28% |
| 22 | 75.04% | 22 | 20.06% |
| 23 | 52.23% | 23 | 22.69% |
| 24 | 62.65% | 24 | 16.28% |
| 25 | 55.26% | 25 | 19.83% |
| 26 | 72.32% | 26 | 2.07% |
| 27 | 91.66% | 27 | 11.62% |
| 28 | 93.05% | 28 | 12.25% |
| 29 | 58.74% | 29 | 7.12% |
| 30 | 76.94% | 30 | 5.97% |
|  |  | 31 | 14.96% |

X% = (negative control average OD_450nm_ value − serum sample OD_450nm_ value)/

(negative control average OD_450nm_ value − positive control average OD_450nm_ value).

Test standard: positive: X% ≥ 50%; negative: X% < 40%; Equivoca: 40% ≤ X% < 50%.

**Supplementary Table 3 Representative ASFV strains of P30’s epitope conservation analysis**

| Strain | Genotype | GenBank accession number |
| --- | --- | --- |
| BA71V | I | KP055815.1 |
| Benin 97/1 | I | NC_044956.1 |
| E75 | I | NC_044958.1 |
| OURT_88/3 | I | NC_044957.1 |
| Pig/HeN/ZZ‑P1/2021 | I | MZ945536.1 |
| Pig/SD/DY‑I/2021 | I | MZ945537.1 |
| Pig/Henan/123014/2022 | I | OQ504954.1 |
| Belgium 2018/1 | II | LR536725.1 |
| Georgia 2007/1 | II | FR682468.2 |
| Korea/pig/Yeoncheon1/2019 | II | MW049116.2 |
| Tanzania/Rukwa/2017/1 | II | LR813622.1 |
| Wuhan 2019‑1 | II | MN393476.1 |
| SPEC-257 | III | DQ250120.1 |
| RSA/99/1/W | IV | EU874307.1 |
| MK-200 | V | MK211506.1 |
| SPEC/265 | VI | EU874264.2 |
| RSA/98/1 | VII | EU874312.2 |
| Malawi/1978 | VIII | JQ744998.1 |
| Ken06.Bus | IX | NC_044946.1 |
| Ken09Tk.13/1 | X | HM745382.1 |
| KAB/62 | XI | EU874289.1 |
| SUM/1411 | XIII | EU874287.1 |
| TAN/01/1 | XV | EU874303.2 |
| TAN/03/2 | XVI | EU874255.2 |
| Zim MVM_90/1 | XVII | JQ745037.1 |
| NAM/P1/1995/C3 | XVIII | PP107957.1 |
| RSA/96/3 | XIX | EU874283.2 |
| RSA/95/1 | XX | EU874266.1 |
| RSA_96/1 | XXI | JQ745031.1 |
| SPEC-245 | XXII | JQ745023.1 |

**Supplementary Table 4 Representative ASFV strains of CD2’s epitope conservation analysis**

| Strain | Genotype | GenBank accession number |
| --- | --- | --- |
| BA71V | I | KP055815.1 |
| Benin 97/1 | I | NC_044956.1 |
| E75 | I | NC_044958.1 |
| NHV | I | NC_044943.1 |
| OURT 88/3 | I | NC_044957.1 |
| K-49 | I | KM609339.1 |
| Pig/HeN/ZZ-P1/2021 | I | MZ945536.1 |
| Pig/SD/DY-I/2021 | I | MZ945537.1 |
| Pig/Henan/123014/2022 | I | OQ504954.1 |
| Belgium 2018/1 | II | LR536725.1 |
| Estonia 2014 | II | LS478113.1 |
| Georgia 2007/1 | II | FR682468.2 |
| Georgia 2008/1 | II | MH910495.1 |
| Korea/pig/Yeoncheon1/2019 | II | MW049116.2 |
| SY-1 | II | OM161110.1 |
| Tanzania/Rukwa/2017/1 | II | LR813622.1 |
| Wuhan 2019-1 | II | MN393476.1 |
| Liv13/33 (OmLF2) | III | MN913970.1 |
| MK-200 | V | KM609347.1 |
| SPEC_57 | VIII | MN394630.3 |
| R35 | VIII | MH025920.1 |
| Ken06.Bus | IX | NC_044946.1 |
| TAN/16/Magu | IX | ON409980.1 |
| Ken05/Tk1 | X | NC_044945.1 |
| Kenya 1950 | X | NC_044944.1 |
| Uvira B53 | X | MT956648.1 |
| TAN/08/Mazimbu | XV | MT956648.1 |
| NAM/P1/1995/C3 | XVIII | PP107957.1 |
| Zaire | XX | MT956648.1 |
| Pretoriuskop/96/4 | XX | AY261363.1 |

**Supplementary Table 5 Representative ASFV strains of P54’s epitope conservation analysis**

| Strain | Genotype | GenBank accession number |
| --- | --- | --- |
| BA71V | I | KP055815.1 |
| Benin 97/1 | I | NC_044956.1 |
| E75 | I | NC_044958.1 |
| OURT_88/3 | I | NC_044957.1 |
| Pig/HeN/ZZ-P1/2021 | I | MZ945536.1 |
| Pig/SD/DY-I/2021 | I | MZ945537.1 |
| Pig/Henan/123014/2022 | I | OQ504954.1 |
| Belgium 2018/1 | II | LR536725.1 |
| Georgia 2007/1 | II | FR682468.2 |
| Korea/pig/Yeoncheon1/2019 | II | MW049116.2 |
| Tanzania/Rukwa/2017/1 | II | LR813622.1 |
| Wuhan 2019-1 | II | MN393476.1 |
| SPEC-257 | III | DQ250120.1 |
| RSA/99/1/W | IV | EU874369.1 |
| MOZ1960/PIR | V | PQ035955.1 |
| SPEC/265 | VI | EU874344.1 |
| RSA/98/1 | VII | EU874374.1 |
| Malawi/1978 | VIII | KC662380.1 |
| Ken09Tk.13/1 | X | HM745325.1 |
| Ken06.Bus | IX | NC_044946.1 |
| KAB/62 | XI | EU874331.1 |
| SUM/1411 | XIII | EU874357.1 |
| TAN/01/1 | XV | EU874356.1 |
| TAN/03/1 | XVI | EU874354.1 |
| ZIM/92/1 | XVII | EU874345.1 |
| NAM/P1/1995/C3 | XVIII | PP107957.1 |
| RSA/96/3 | XIX | EU874375.1 |
| RSA/95/1 | XX | EU874340.1 |
| RSA/96/1 | XXI | EU874339.1 |
| ETH/1a | XXIII | KT795363.1 |

**Supplementary Table 6 Representative ASFV strains of P72’s epitope conservation analysis**

| Strain | Genotype | GenBank accession number |
| --- | --- | --- |
| BA71V | I | KP055815.1 |
| Benin 97/1 | I | NC_044956.1 |
| E75 | I | NC_044958.1 |
| NHV | I | NC_044943.1 |
| OURT 88/3 | I | NC_044957.1 |
| Pig/HeN/ZZ-P1/2021 | I | MZ945536.1 |
| Pig/SD/DY-I/2021 | I | MZ945537.1 |
| Pig/Henan/123014/2022 | I | OQ504954.1 |
| Belgium 2018/1 | II | LR536725.1 |
| Georgia 2007/1 | II | FR682468.2 |
| Korea/pig/Yeoncheon1/2019 | II | MW049116.2 |
| Tanzania/Rukwa/2017/1 | II | LR813622.1 |
| Wuhan 2019-1 | II | MN393476.1 |
| Liv13/33 (OmLF2) | III | MN913970.1 |
| Warthog | IV | NC_044949.1 |
| Tengani 62 | V | AY261364.1 |
| Mkuzi 1979 | VII | AY261364.1 |
| R35 | VIII | MH025920.1 |
| SPEC_57 | VIII | MN394630.3 |
| Ken06.Bus | IX | NC_044946.1 |
| TAN/16/Magu | IX | ON409980.1 |
| BUR/18/Rutana | X | MW856067.1 |
| Ken05/Tk1 | X | NC_044945.1 |
| Kenya 1950 | X | NC_044944.1 |
| Uvira B53 | X | MT956648.1 |
| TAN/08/Mazimbu | XV | MT956648.1 |
| NAM/P1/1995/C3 | XVIII | MT956648.1 |
| Zaire | XX | MT956648.1 |
| Pretoriuskop/96/4 | XX | AY261363.1 |
| ETH/1a | XXIII | KT795359.1 |

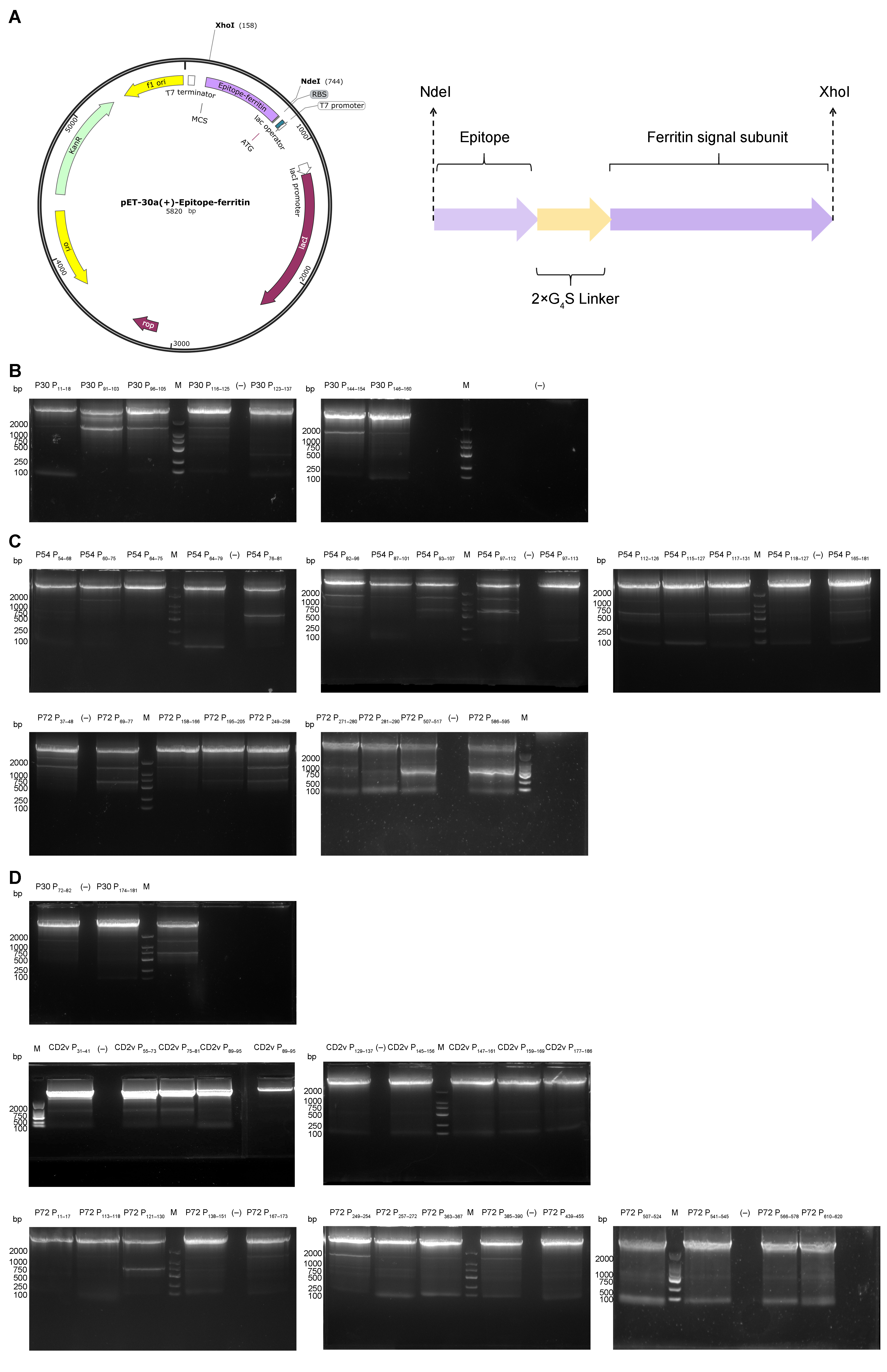

**Supplementary Fig. 1 Construction and verification of the pET-30a(+)-epitope-ferritin expression plasmid. (A)** Map of the pET-30a(+)-epitope-ferritin plasmid. Key genetic elements include the Kanamycin resistance gene, the T7 promoter, and the inserted gene fusion encoding the epitope fused to the Pyrococcus furiosus ferritin signal subunit (as shown in the map, specifically CD2v P31–41). The fusion gene was cloned between the NdeI and XhoI restriction sites. **(B)** PCR analysis of pET-30a(+)-P30 (IEDB)-epitope-ferritin. Product size: ~5,000 bp. M: marker; (–): negative control. **(C)** PCR analysis of pET-30a(+)-P54 (IEDB)- and P72 (IEDB)-epitope-ferritin plasmids. Products: ~5,000 bp. M: marker; (–): negative control. **(D)** PCR analysis of pET-30a(+)-P30 (Prediction)- CD2v (Prediction and IEDB)-, and P72 (Prediction)-epitope-ferritin plasmids. Products: ~5,000 bp. M: marker; (–): negative control.

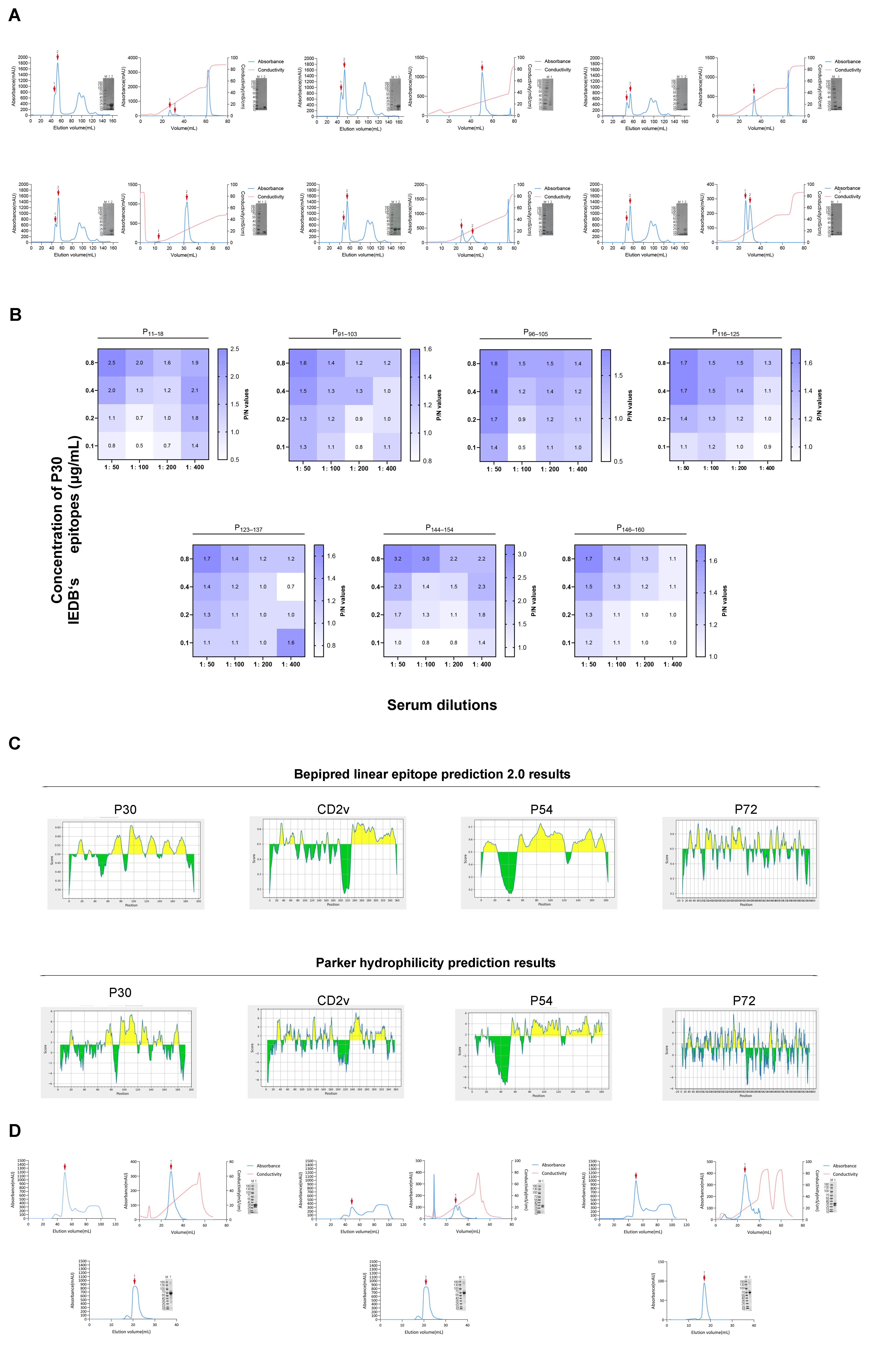

**Supplementary Fig. 2 P30’s EFNs’, GST-His_6_-epitops’ and BSA-conjugated eotopes’ purification. ELISA optimization and prediction method comparison. (A)** Purification of P30 P_91–103_–EFNs, P_96–105_–EFNs, P_116–125_–EFNs, P_123–137_–EFNs, P_144–154_–EFNs, P_146–160_–EFNs. Left: size-exclusion chromatography profile with major peaks (red arrows) corresponding to 24-mer nanoparticle complexes (~450 kDa). Inset: SDS-PAGE analysis confirming monomeric ferritin (~19 kDa). Right: anion exchange chromatography profile with major peaks (red arrows) corresponding to 24-mer nanoparticle complexes (~450 kDa). Inset: SDS-PAGE analysis confirming monomeric ferritin (~19 kDa). **(B)** Optimization of EFNs-iELISA conditions by checkerboard titration method. **(C)** Length evaluation of predicted linear B-cell epitopes between Bepipred Linear Epitope Prediction 2.0and Parker Hydrophilicity Prediction. **(D)** Purification of GST-His₆–tagged and BSA-conjugated peptides containing P30 epitopes P_11–18_, P_96–105_ and P_144–145_.

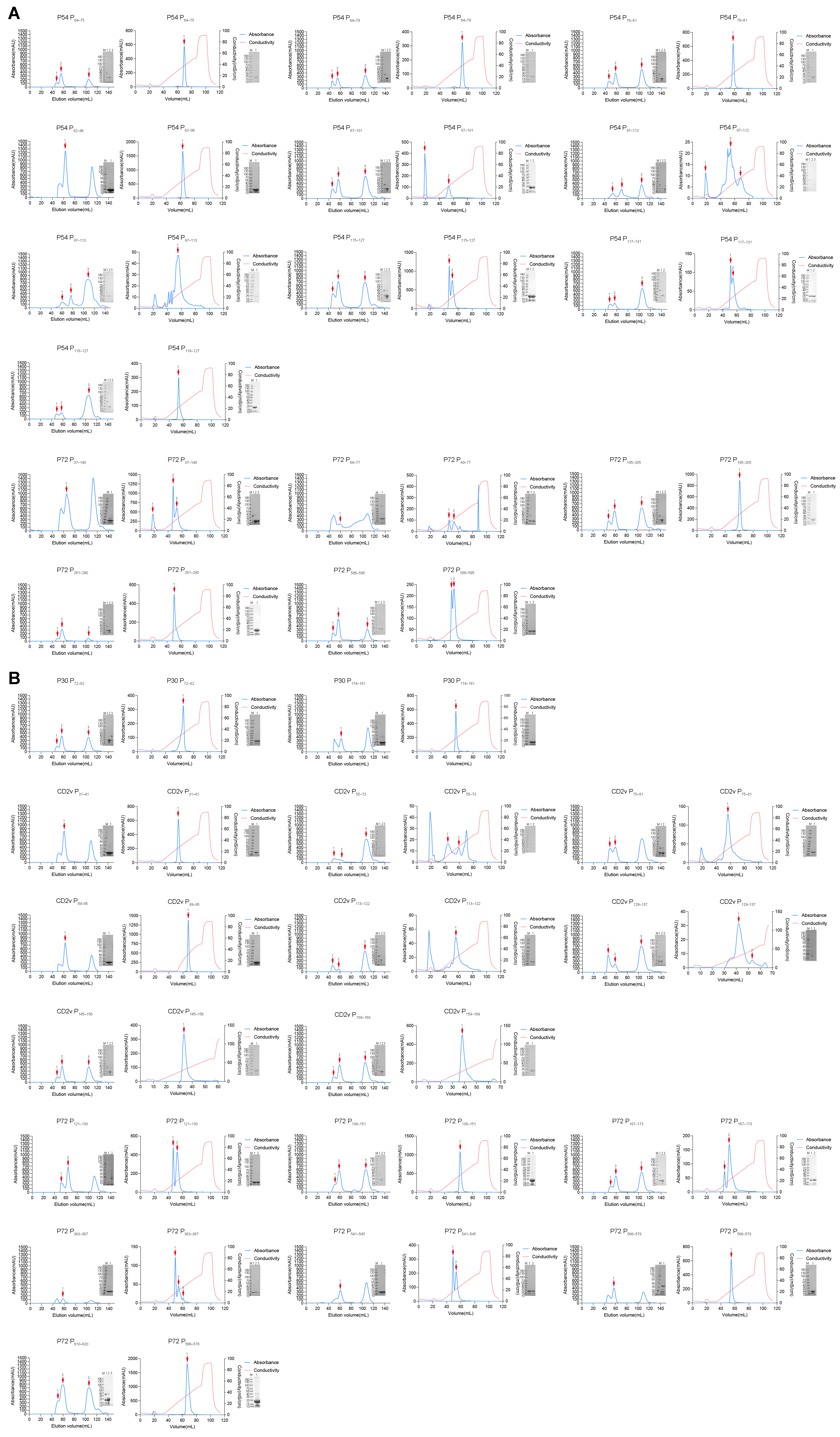

**Supplementary Fig. 3 CD2v, P30, P54, and P72’s EFNs purification. (A)** Purification of P54 and P72’s EFNs from IEDB. **(B)** Purification of P30, CD2v, and P72’s EFNs from prediction.
